# Antibiotic resistance genes in agriculture and urban influenced watersheds in southwestern British Columbia

**DOI:** 10.1101/104851

**Authors:** Miguel I. Uyaguari-Díaz, Matthew A. Croxen, Zhiyao Luo, Kirby I. Cronin, Michael Chan, Waren N. Baticados, Matthew J. Nesbitt, Shaorong Li, Kristi Miller, Damion Dooley, William Hsiao, Judith L. Isaac-Renton, Patrick Tang, Natalie Prystajecky

**Author notes:** Correspondence: Miguel I. Uyaguari-Díaz, British Columbia Centre for Disease Control Public Health Laboratory, Vancouver, British Columbia, Canada V5Z 4R4.

## Abstract

**Background:** The dissemination of antibiotic resistance genes (ARGs) from anthropogenic activities into the environment poses an emerging public health threat. Water constitutes a major vehicle for transport of both biological material and chemical substances. The present study focused on putative antibiotic resistance and integrase genes present in the microbiome of agricultural, urban influenced and protected watersheds in southwestern British Columbia, Canada. A metagenomics approach and high throughput quantitative PCR (HT qPCR) were used to screen for elements of resistance including ARGs and integron-associated integrase genes (*intI*). Sequencing of bacterial genomic DNA was used to characterize the resistome of microbial communities present in watersheds over a one-year period.

**Results:** Data mining using CARD and Integrall databases enabled the identification of putative antibiotic resistance genes present in watershed samples. Antibiotic resistance genes presence in samples from various watershed locations was low relative to the microbial population (<1 %). Analysis of the metagenomic sequences detected a total of 78 ARGs and *intI1* across all watershed locations. The relative abundance and richness of antibiotic resistance genes was found to be highest in agriculture impacted watersheds compared to protected and urban watersheds. Gene copy numbers (GCNs) from a subset of 21 different elements of antibiotic resistance were further estimated using HT qPCR. Most GCNs of ARGs were found to be variable over time. A downstream transport pattern was observed in the impacted watersheds (urban and agricultural) during dry months. Urban and agriculture impacted sites had a higher GCNs of ARGs compared to protected sites. Similar to other reports, this study found a strong association between *intI1* and ARGs (e.g., *sul1*), an association which may be used as a proxy for anthropogenic activities. Chemical analysis of water samples for three major groups of antibiotics was negative. However, the high richness and GCNs of ARGs in impacted sites suggest effects of effluents on microbial communities are occurring even at low concentrations of antimicrobials in the water column.

**Conclusion:** Antibiotic resistance and integrase genes in a year-long metagenomic study showed that ARGs were driven mainly by environmental factors from anthropogenized sites in agriculture and urban watersheds. Environmental factors accounted for almost 40% of the variability observed in watershed locations.

## Introduction

Antibiotic resistance is recognized as a major emerging health threat worldwide. While not a new phenomenon, factors such as the increasing global population growth, overuse of antibiotics, and limited development of new drugs have increased the risks of morbidity and mortality as well as the costs associated with the treatment of bacterial diseases [1, 2].

Globally, the annual consumption of antibiotics is estimated to be 70 billion standard units/year for human use [3] and 63,151 ± 1,560 tons/year for livestock [4]. Within the next 15 years, usage is expected to increase by 30 % and 67 % for human and veterinary purposes, respectively [5]. Currently, mortality rates attributable to antimicrobial resistance represents 700,000 deaths annually [6]. Some reports estimate this rate will increase to 10 million deaths/year by 2050, with antimicrobial resistance becoming the leading cause of death world-wide [6]. While this estimate is being debated [7], an exponential trend will undoubtedly continue as a consequence of the use and misuse of antibiotics associated with clinical and agricultural use.

It is estimated that between 75 % and 90 % of antibiotics are poorly absorbed by either humans or animal hosts, and excreted unaltered in feces or urine [8-10]. Thus spillage of antibiotics and their metabolites into the environment is widespread with contamination hotspots including hospital sewage discharges, health care facility and community wastewater treatment plants (WWTPs), pharmaceutical industry facilities, and confined animal feeding facilities [11, 12]. Impacted terrestrial and aquatic ecosystems such as soil, rivers, streams, watersheds, groundwater and sediments, are the recipients of both antibiotic residues and antibiotic-resistant bacteria [12, 13]. While some antibiotics may degrade quickly, others accumulate in soil or sediments and remain for a longer period of time [14, 15]. Surface water remains the main vehicle of dissemination of these residues, antibiotic resistant bacteria and antibiotic resistance gene elements into the environment [16, 17].

Microbial acquisition of genes involved in antibiotic resistance mechanisms is a result of a variety of gene transfer systems or elements such as conjugative plasmids, transposons, and integrons [18, 19]. Regardless of the gene donor, these horizontal gene transfer elements enable genes to move from one bacterial cell to another and from one genomic system to another [18]. The role of elements such as phages and integrons in the spread of resistance appears to be significant in freshwater ecosystems [20] and has been well documented using culture-independent approaches [21-24]. In this context, several metagenomic studies have been conducted in aquatic ecosystems analyzing the resistome of microbial communities from sediments, WWTPs and effluents [25-28]. Only a few, however, have focused on antibiotic resistance genes or elements of resistance in surface water or watersheds [29, 30]. Watersheds are the primary source of drinking water and represent a pivotal route affected by land-use and anthropogenic activities flowing into other larger aquatic ecosystems such as rivers, basins and seas. These water bodies likely contain antibiotic residues or metabolites, and thus microorganisms in watersheds may carry antibiotic resistance genes that play a key role in the emergence of antibiotic resistance in human pathogens. In the present study, we collected raw surface water samples from 3 watersheds in southwestern British Columbia, Canada over a one-year period. Each watershed was characterized by different land-use (agricultural, urban and protected sites). Sequence-based metagenomics and high-throughput quantitative PCR were used to characterize and quantify elements of resistance in these water samples. Understanding the interplay between land use, watersheds and the dissemination of antibiotic resistance genes or genetic elements associated to resistance may help avert future impacts to public health and to the environment.

## Results and Discussion

Water samples from seven sampling sites located in three watersheds of differing land-use in southwestern British Columbia were collected over the course of one year. Bacterial metagenomics generated 36 Gb for this dataset. CARD and Integrall databases were selected over other classification methods due to underestimation (ARG-annot and VFDB) and overestimation (SEED subsystems) of the other databases (Additional file 1: Table S1). A total of 3.95 million contigs (≥ 200 bp) were assembled and analyzed using CARD and Integrall. CARD identified 12,044 hits from 6,640 contigs as elements of resistance with grouping into 78 unique antibiotic resistance gene types conferring resistance to at least one antibiotic. Integrall database identified 6,711 hits with 149 contigs associated to *intI1.*

### Main classes of antibiotics and mechanisms of antibiotic resistance in **watersheds**

CARD identified a large proportion (~90 %) of the contigs as housekeeping function-associated genes such as elongation factors (55.4 %), DNA-directed RNA polymerase (23.8 %) and DNA topoisomerases/gyrases (10.6 %) (Additional file 2: Figure S1a). Although these functions can be targeted by antibiotics such as elfamycin, daptomycin/rifampin and fluoroquinolones (Additional file 3: Figure S1b), respectively, the identification of these genes is not an indication of resistance; the universality of these structural genes has been widely documented in a variety of environments [26, 31-33]. A small percentage (~10 %) of genes was found to encode resistance to at least one antibiotic through a protein or were involved in multidrug resistance/efflux systems. Within this group, genes involved in multiple drug resistance (MDR) made up at least ~39 % of transport of efflux pumps in the protected downstream site (PDS) (Fig. 1a). This proportion is in agreement with previous estimates of bacterial genes involved in mechanisms of transport, and mainly genes encoding efflux pumps [34]. In this context, major mechanisms of efflux pumps included: resistance-nodulation-division ranging from 5 % (PDS) to 51 % (protected upstream, PUP); multidrug efflux system proteins with values between 5 % and 30 % in PDS and urban polluted (UPL), respectively; and genes associated with transport at <5 % across all sites (Fig. 1b).

**Figure 1.**
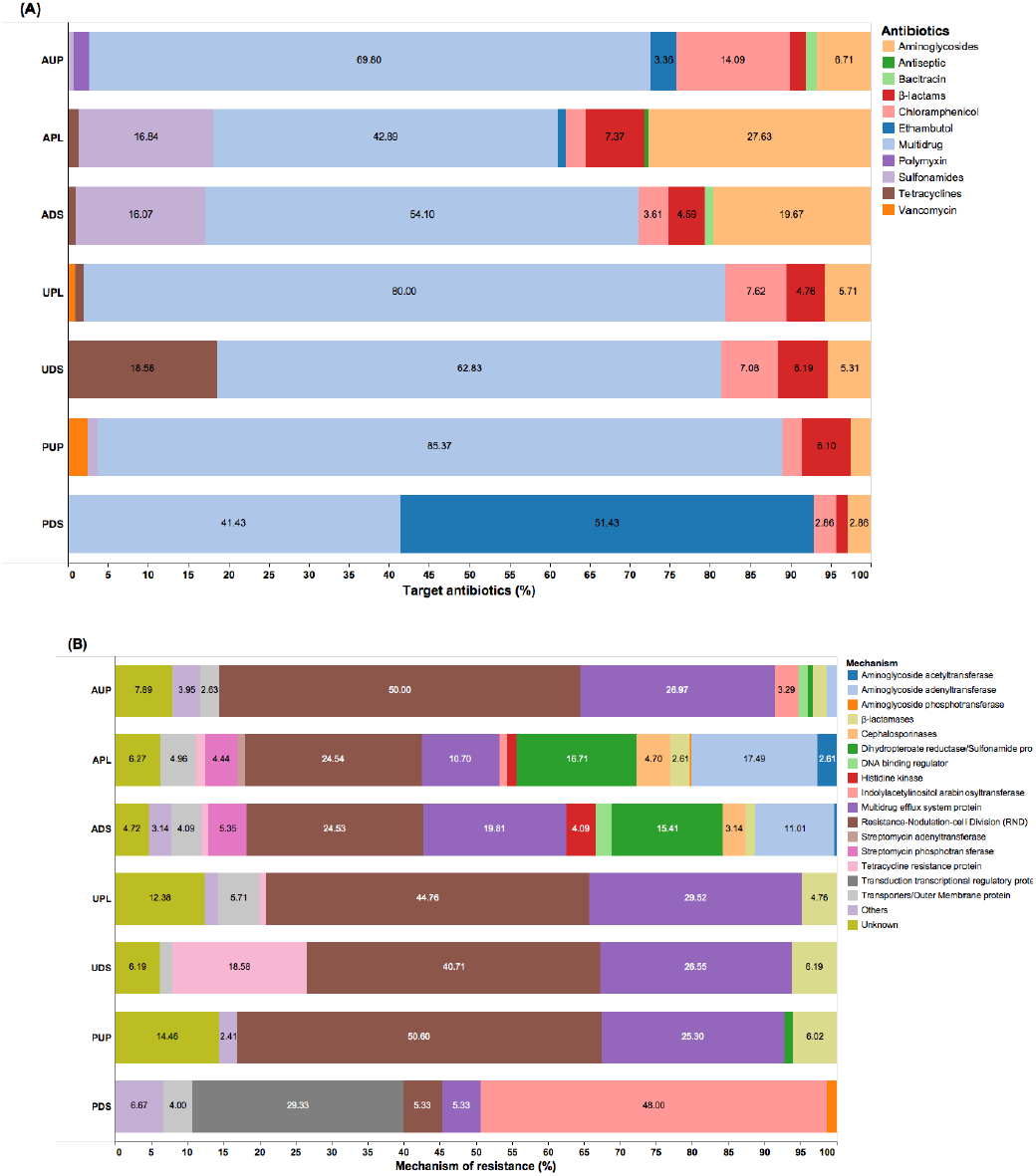
Composition of antibiotic resistance gene categories assigned by CARD in watershed locations: A) Group of antibiotic, and B) Mechanism of action. AUP: agricultural upstream site; APL: agricultural polluted; ADS: agricultural downstream; UPL: urban polluted; UDS: urban downstream; PUP: protected upstream; PDS: protected downstream.

In the agriculture impacted watershed study sites, relative abundance of ARGs affiliated with aminoglycosides (agricultural polluted, APL: 27.6 % and agricultural downstream, ADS: 19.7 %) and sulfonamides (APL: 16.8 % and ADS, 16.1 %) were high compared to the urban and protected sampling sites. Moreover, abundance of genes associated to chloramphenicol/phenicol resistance groups observed in the urban influenced sites were at least 3 fold higher than agricultural or protected sites (Fig. 1a). Although agricultural upstream watershed sampling site (AUP), is located upstream of agricultural activity, it should be noted that there is low density residential land-use in this area and may have had an effect on the relative abundance of ARGs related to chloramphenicol/phenicols. Moreover, genes conferring resistance to tetracyclines were observed in both the agricultural and urban impacted watersheds. A high relative abundance of these genes was observed in urban downstream or UDS (18.6 %). We hypothesize that this elevated value may be due to water from the UPL site traveling 1.1 km through parkland and some area with urban development (non-point source contamination) before merging into the UDS site. Except for AUP and PDS, no major differences were observed across watersheds for ARGs encoding for β-lactamases (Fig. 1a and 1b). In protected sites, genes associated to vancomycin and ethambutol resistance made up 2.4 % and 51.4 % of the resistome in PUP and PDS, respectively. Given the occurrence of these genes in protected sites, these values may be related to naturally occurring bacteria harboring component system regulators such as *embB*-*embC*, and *vanR*-*vanS*. Other gene groups associated with resistance to bacitracin, polymyxin, and regulatory component systems of heavy metals such as copper, iron-sulfur and tellurite ions, were observed in agricultural sites and PDS. These results confirm the prevalence of antibiotic resistance determinants to these major classes of antibiotics (β-lactams, phenicols, aminoglycosides, and tetracyclines) in anthropogenic-influenced environments [25, 26, 35, 36].

### Diversity and distribution of antibiotic resistance genes in watersheds

From the bacterial DNA metagenomes, CARD and Integrall database identified a total of 78 ARGs and one *intI1* gene associated with the different watersheds. Figure 2 shows the distribution of ARGs and *intI* in the watershed locations. A total of 68 out of 79 genes (including *intI*) were observed in agriculture influenced watersheds. A large number of these ARGs were observed in APL (n=38) and ADS (n=29), while less richness of ARGs genes was found in AUP (n=11) (Fig. 2a). In AUP, ARGs with high relative abundance included *mdtB* (17.6 %), *acrD* (11.8 %), *mexB* (17.6 %), and *mexY* (14.7 %). In APL, *intI1* made up a large proportion of the elements of resistance (34.7 %) (Fig. 2a). Other representative ARGs in APL included *sul1* (11.6 %), *aadA1* (6.7 %), *mexY* (5.9 %), *mexB* and *ampC* with 5.0 % each, and *sul2* (2.0 %). In ADS, *sul1* (21.2 %) was the ARG with high relative abundance, while the *intI* genes could not be detected using a metagenomics approach. These findings are consistent with previous studies reporting prevalence of *sul1* and *sul* genes in agriculture impacted environments [37, 38]. Other abundant genes in ADS included *mexB* (7.0 %) and *mexY* (8.5 %), *aadA1* (6.1 %) and *strB* (6.6 %). Overall, we observed a wide range of aminoglycosides resistance genes in both agriculture impacted sites, APL (n=20) and ADS (n=18) (Fig. 2a). Although detected in low relative ratios, *tet (W)* was found in agriculture impacted watersheds (APL and ADS) with average values of 1.3 %. Groups such as *intI* and *sul* genes were prevalent in both APL and ADS.

**Figure 2.**
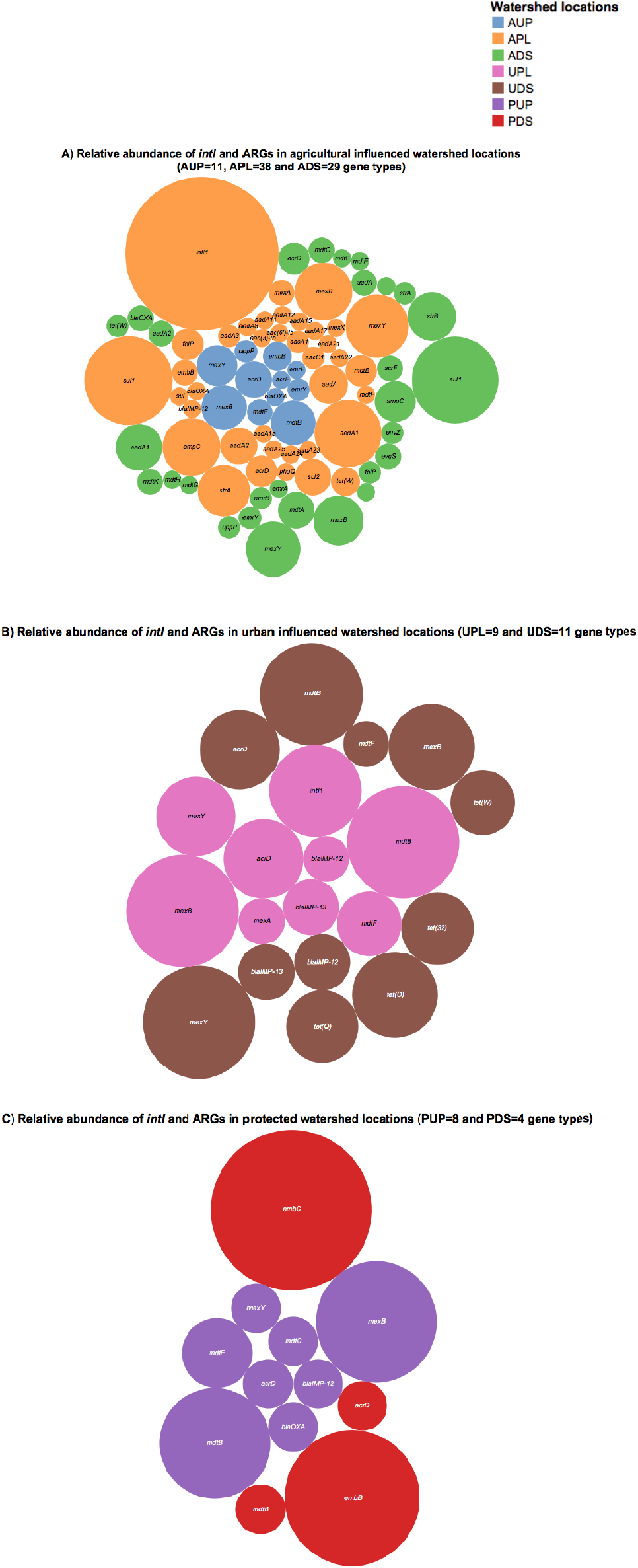
Relative abundance of integrase and antibiotic resistace genes in A) agricultural, B) urban DNA, and C) protected watershed locations. AUP: agricultural upstream site; APL: agricultural polluted; ADS: agricultural downstream; UPL: urban polluted; UDS: urban downstream; PUP: protected upstream; PDS: protected downstream.

Urban sites were less diverse in ARGs compared to APL and ADS. In UPL and UDS, a total of 9 and 11 ARGs were observed, respectively. From the sequencing data, we observed that urban sites shared at least 8 ARGs between them. On average, genes such as *mexB* (17.9 %)*, mdtF* (4.9 %), *mexB* (15.7 %) and *mexY* (14.3 %) made up the majority of ARGs in UPL and UDS (Fig. 2b). Moreover, *intI* genes were detected in different proportions in UPL (13.8 %) and UDS (1.5 %). Similar to the agriculture impacted sites, UPL had a high relative abundance of *intI1*, which may be an indication of the prevalence of these genes in anthropogenic impacted environments [22, 39, 40]. Moreover, tetracycline resistance genes were identified in UDS and UPL watersheds: *tet(32)* (7.6 %), *tet (O)* (10.6 %), *tet (Q)* (7.6 %), *tet (W)* (6.0 %), and *otrC* (1.72 %) (Fig. 2b). More generally, in urban sites we observed determinants of antibiotic resistance such as *intI*, *sul*, and *tet* genes, which have been reported to be associated with wastewater and surface water receiving such effluents [41, 42].

When compared to agricultural and urban locations, a low diversity of ARGs was observed in protected watersheds. A total of 10 types of ARGs were found in protected watersheds, and distributed into 8 and 4 types for PUP and PDS, respectively. In PUP, *mdtB* and *mexB* made 25 % and 30 % of ARGs, while in PDS, *embB* and *embC* comprised 85.7 % of the ARGs (Fig. 2c). Integrase genes were not detected in any of these watershed locations using a metagenomics approach. Genes such *acrD* and *mdtB* were observed in both sites with average values of 5 % and 7.5 %, respectively. The diversity of β-lactamases such as *bla*_LRA_, *bla*_IMP_, and *bla*_OXA_ was observed in small percentages ranging from 2.4 to 5.0 % even in the protected watershed sites. This is consistent with reports where carbapenemase genes have been detected in remote areas without human activity and is thought to be associated with pathogenic organisms harboring these genes [43]. Similar to UPL, *vanRS*, a two component regulatory system was detected in PUP. The presence of this system may be associated with members of *Nocardiaceae* or other Gram-positive bacteria present in soil [44, 45]. Although phylogenetic information is associated with most of these genes in the databases, it is difficult to assign a particular genus or family to a particular gene. Thus, the information described here focused on the elements of resistance. Overall, agriculture impacted sites were found to have greater richness of ARGs compared to urban and protected locations. Our metagenomics approach detected *intl1*, a potential indicator of gene cassettes in only the impacted sites, while *intI2* nor *intI3* were detected in of the study sites. To quantify these elements of resistance in watersheds further, we complemented the information generated from the bacterial metagenomics study with high throughput qPCR (HT qPCR) analysis.

### Detection and quantitation of antibiotic resistance genes in watershed **locations**

To confirm and quantify ARG prevalence in the study watersheds, primer and probes were designed for 60 elements of resistance, based on results from the metagenomics study. Although only class 1 of *intI* genes was detected by metagenomics, we incorporated two other main classes of integrase genes (*intI2* and *intI3*) that have been reported in the literature [46]. Samples collected from the same locations, but not part of this study, were used to validate qPCR assays for the ARGs. The primers were first checked for sensitivity and specificity by SYBR green-based PCR (data not shown). TaqMan probes (Life Technologies, Carlsbad, CA) were then designed and validated for the primers which showed suitable sensitivity and specificity (data not shown). Assays with cycle threshold values greater than 35 were not included in the HT qPCR run using the BioMark system (Fluidigm Corporation, South San Francisco, CA). The final qPCR panel included a total of 18 ARG, 3 classes of *intI*, and 16S rRNA gene primers and probes (Additional file 1: Table S2>) which were run on DNA templates from the different watershed study sites. A heat map of antibiotic resistance genes normalized by bacterial counts (as estimated by the 16S rRNA gene) shows the relative abundance and distribution of each gene in watershed locations over time (Fig. 3). Most of the ARGs were estimated to be present in an order of magnitude of 1 × 10^−2^ (Fig. 3). These findings are in agreement with other studies conducted in agriculture and urban influenced environments [21, 29, 47]. Genes such as *mdtF*, *soxR*, *tet(32)*, *tet (W)* had ratios of up to 3.8 × 10^−1^ were found in agriculture impacted sites and UDS. We hypothesize that these values are related to agricultural discharges and in the UDS site, runoff from storm water could explain the high ratio of tetracycline resistance genes. These gene ratios were also high in agriculture impacted sites. When ratios were compared across study sites, genes with more distinctive pattern were observed. For instance, genes such as *aacA1*, *aadA1*, *strA*, *strB*, *sul1*, *sul2* showed high ratios in the agriculture impacted sites, while relative concentrations as high as 3.2 × 10^−1^ and 1.5 × 10^−1^ were observed for *tet(32)* and *tet (Q)*, respectively in the urban impacted watersheds. Gene *tet (W)* was prevalent in both urban and agriculture impacted sites compared to protected watersheds. We also observed that the *intI1* gene was present over the whole year and in all sites (April 2012-April 2013). Class 1 integrase genes had ratios ranging from 3.4 to 3.7 × 10^−2^ in ADS and APL, respectively, while those for AUP had a ~5.8 × 10^−3^ relative concentration. Ratios detected in agriculture impacted sites were relatively high compared to urban impacted (1.1-1.9 × 10^−2^) and protected watersheds (1.6-7.7 × 10^−3^). Class 2 integrase genes were detected in agricultural sites (1.6-1.9 × 10^−3^) and to a lesser degree in urban sites ranging from 7.0 × 10^−5^ (UPL) to 1.4 × 10^−4^ (UDS), while in protected sites, *intI2* was only detected once in PUP, at a low relative concentration (8.0 × 10^−5^). Although intermittently observed in the protected watershed locations, integrase class 3 was detected in all watershed locations. On average concentrations of *intI3* were one order of magnitude higher in downstream sites of impacted watersheds (ADS and UDS) compared to the other sites. Some gene patterns in terms of seasonality were observed between the dry (time points 1-7, corresponding to spring and summer) and rainy season (time points 8-13, corresponding to fall and winter) (Fig. 3); more time series studies across seasons are needed to confirm this observation. The prevalence of certain elements of resistance over time may serve as proxies of anthropogenic impacts, as proposed by other studies [48-50]. In this context and to elucidate differences over time of ARGs and *intI* in watershed sites, we conducted repeated measures analysis in terms of sample volume and total extracted bacterial biomass.

**Figure 3.**
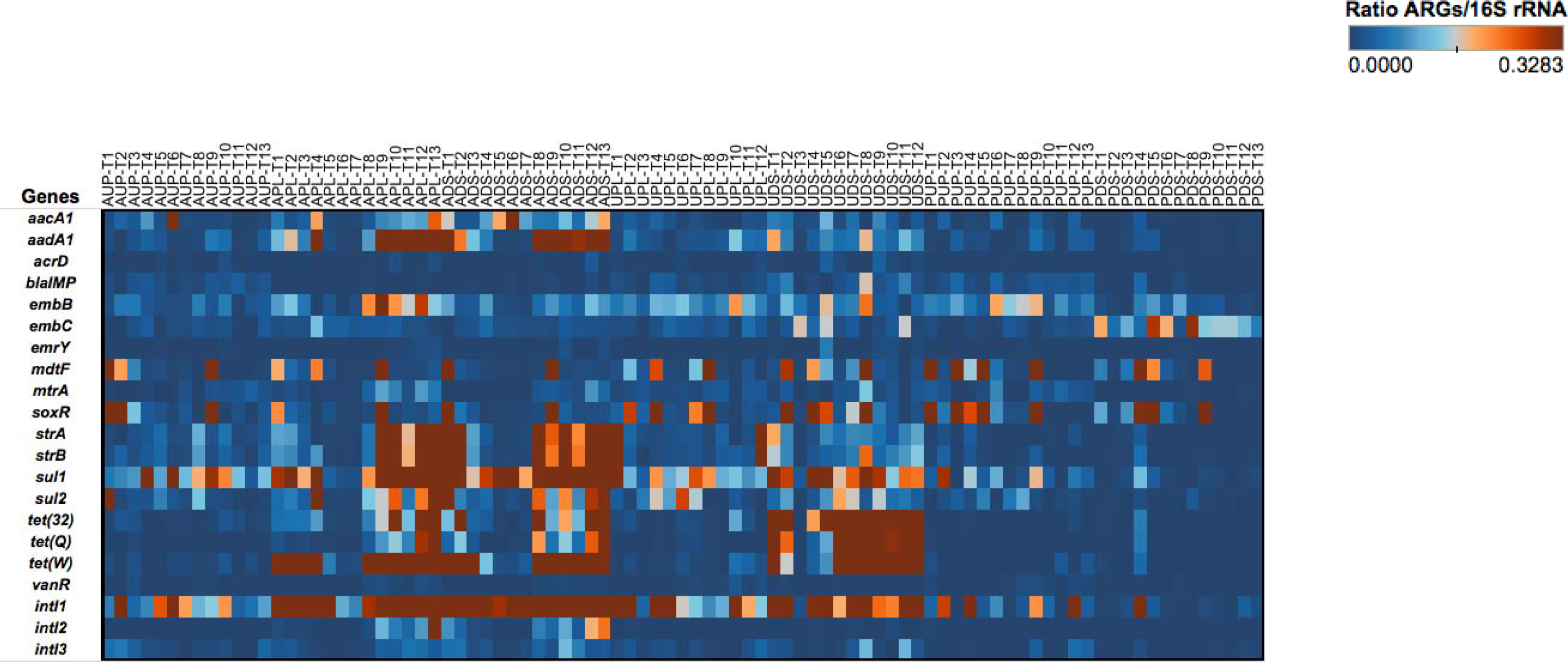
Heat map depicting the ratio of antibiotic resistance genes and 16S rRNA gene over time (as estimated by high-throughput quantitative PCR platform) in watershed sites. AUP: agricultural upstream site; APL: agricultural polluted; ADS: agricultural downstream; UPL: urban polluted; UDS: urban downstream; PUP: protected upstream; PDS: protected downstream. Months are represented by T1 through T13 that corresponding to the long-year study (April 2012- April 2013).

### Repeated measures analysis in watershed locations

Absolute numbers generated from standard curves in the HT qPCR platform were normalized per ml of sample (volume) and ng of water DNA (biomass) as described by Ritalahti et al. [51]. Due to the multicopy nature of the 16S rRNA gene, a factor of 4.3 was used to normalize bacterial counts [52]. Figures 4 and S2 (Additional file 4) depict gene copy numbers per ml of water sample and ng DNA, respectively, for all watershed study sites. Repeated measures analysis revealed striking differences between watershed sites in terms of 16S rRNA gene, ARGs and *intI* genes. Copy numbers of the 16S rRNA gene in orders of magnitude of 10^5^ (volume) were detected and found to change over time within the watersheds. When comparing sites, we observed average quantities of the 16S rRNA gene with 7.34 × 10^5^ GCN/ml sample in PDS, followed by ADS and APL locations with 5.67 × 10^5^ and 4.17 × 10^5^ GCN/ml of sample, respectively. It is probable that values observed in agriculture impacted watersheds are associated with farm discharges. We propose that the high GCN in PDS may be due to biofilms in the pipe where study samples were collected [53].

**Figure 4.**
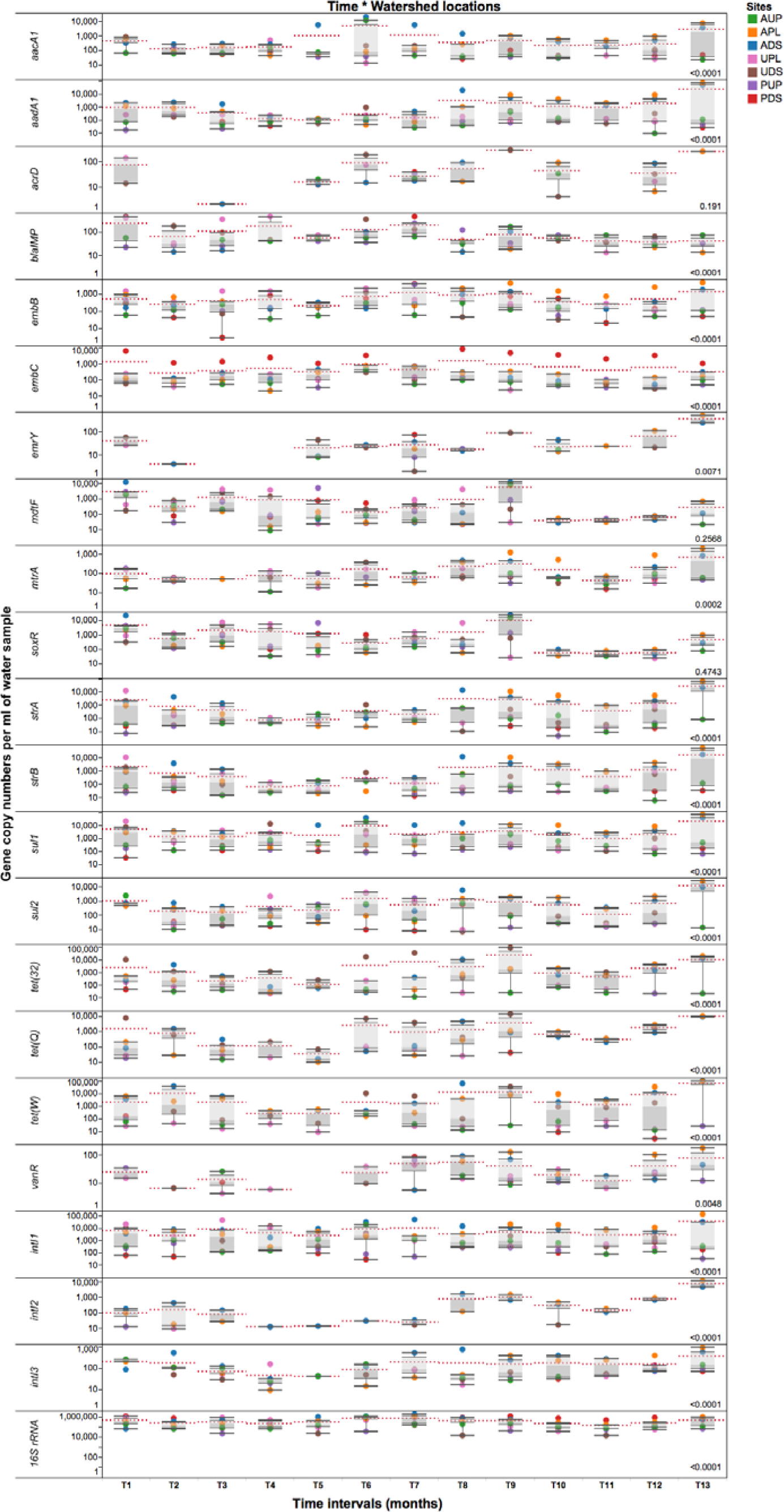
Absolute gene copy numbers of antibiotic resistance genes, integrase gene classes 1, 2, and 3, and 16S rRNA gene per ml of environmental water sample over time in the different locations. AUP: agricultural upstream site; APL: agricultural polluted; ADS: agricultural downstream; UPL: urban polluted; UDS: urban downstream; PUP: protected upstream; PDS: protected downstream. Red dotted lines represent means for a specific time point. Number on the lower right represents p-value from the repeated measures analysis. Statistical significance was set at the 0.05 level.

Repeated measures analysis did not show significant differences (p>0.05) for GCNs of *acrD*, *mdtF* and *soxR* over time (Fig. 4). In addition to these genes and when normalized per ng of DNA, means of *embC*, and *emrY* were not found to be significantly different (p≥0.5996) over time in the study sites (Additional file 4: Figure S2). Since the GCNs per volume for these two genes were 1 to 2 orders of magnitude higher than GCNs per ng of DNA water, we suggest that the significant decrease of organisms harboring these genes made any difference undetectable over time.

In the agriculture impacted sites, significantly higher GCNs of *aacA1*, *aadA1*, *embB*, *emrY*, *mtrA*, *strA*, *strB*, *sul1*, *sul2*, *intI* classes 1-3 were detected over time compared to the protected watersheds (Fig. 4). Genes such as *emrY*, *tet(32)*, *tet (Q)*, *tet (W)* were more abundant in both the agriculture and urban impacted sites compared to the protected and upstream sites. During this yearlong study, integrase classes 1, 2 and 3 appeared to be predominant in APL and ADS. Moreover, the means of *intI* genes differed (p<0.0001) over time per watershed location. In agricultural sites, *intI* class 1, 2 and 3 genes were detected in at least one order of magnitude higher than in urban or protected watersheds, with values ranging from 10^4^-10^5^, 10^2^-10^4^, and 10^2^-10^3^ GCNs/ml of sample for *intI1*, *2* and *3*, respectively. Integrase class 1 genes were detected in all sites, GCNs were lower in the protected watersheds compared to the agriculture and urban influenced watersheds. Integrase class 2 was mostly observed in APL and ADS over time, while that *intI3* was detected in higher GCNs in agricultural and urban sites. It was observed that in some sampling events *intI2* (T1) or *intI3* (T10-T12) were detected in PDS, but in significantly lower GCNs compared to the impacted sites. We observed that most of these significant changes in absolute GCNs per ml of water occurred around T7 (October), which is the beginning of the local rainy season. For instance, higher GCNs of *bla*_IMP_ were observed in urban influenced watersheds from T1 to T7 (Fig. 4). A slight decline in GCNs of *bla*_IMP_ was detected and persisted over time once the rainy season started (T8-T13). Although AUP is not directly impacted by agriculture activities, the presence of some residences near the sampling site may have some influence in the *bla*_IMP_ gene patterns observed. Moreover, *vanR* was detected intermittently in urban impacted sites during the T1-T6 period (Fig. 4), while after T7, GCNs of this gene increased significantly in agriculture impacted sites. It has been shown that *vanR*, a regulatory structural gene that could also be part of a mobile *van* operon [54], may be carried by *Rhodococcus equii* and *Enterococcus* spp. [45, 55], bacteria commonly found in soil and able to cause pneumonia in grazing animals [56]. These bacteria also may reside in the human gastrointestinal tract, with a potential for causing urinary tract infections, bacteremia and endocarditis in some circumstances [55]. It is also possible that GCNs of *vanR* reflects farm facility discharges or sewage infrastructure seepage in surrounding urban site areas.

An analysis of GCNs expressed per ng DNA (Additional file 4: Figure S2) revealed seasonal downstream transport of ARGs in the sampling sites. We observed these intermittent changes occurred at different times of the year. From T1 through T8 (April-November 2012), high GCNs of *aacA1*, *sul1*, *sul2*, *intI1* and *intI2* were detected in ADS compared to APL (Additional file 4: Figure S2). During the same time period (T1-T8), high GCNs per ng DNA of *bla (IMP)*, *tet(32)*, and *tet (Q)* were observed in UDS compared to UPL (Additional file 4: Figure S2). Moreover, a downstream transport of genes such as *aadA1*, *strA*, *strB* and *tet (W)* was observed in both agricultural and urban downstream locations (Additional file 4: Figure S2). Overall, low GCNs of ARGs were detected during T4 and T5 in all watershed locations (Fig. 4 and Additional file 4: Figure S2), the driest months (July and August) of the year in southwestern British Columbia. This pattern of seasonal variation was similar for most ARGs described (Additional file 4: Figure S2). After T8 and with the onset of the rainy season, higher GCNs were observed. No transport patterns, however, were observed downstream from the impacted sites. Seasonal changes have been documented in other freshwater ecosystems studies [57].

### Exploratory factor analysis of watershed microbiomes

An exploratory factor analysis was conducted using the ratios of antibiotic resistance genes to the 16S rRNA gene, and with physicochemical and biological parameters from each study watershed. Both orthogonal and oblique rotations were conducted with the data. Figure 5 depicts an oblique parsimax rotation that best fits all variables assessed in this study. Factor 1 or “anthropogenic stressors” accounted for 38.7 % of the observed variability, while that factor 2 or “aerobic conditions” accounted for 37.4 %. Four clusters can visually be identified (Fig. 5): APL and ADS; AUP and UPL; UDS; PUP; and PDS. Integrase genes class 1, *sul1*, *sul2*, *strA*, *strB*, *aacA1*, *aadA1*, *tet (W)*, and *embB* seemed to be driven by anthropogenic stressors in agriculture impacted sites (APL and ADS), UDS and to a lesser extent UPL. Moreover, land-use nutrients such as phosphate, nitrate, nitrite and ammonia [58-60] were observed within the same quadrant as these ARGs. In additional models, where metadata were excluded from the analysis or where only nutrients associated to land-use were incorporated into the model, *intI1* and *sul1* clustered within the same quadrant as anthropogenic perturbations (data not shown). Furthermore, a significant positive correlation (r=0.8078, p<0.0001) was detected between these two genes (Additional file 5: Figure S3). This finding is in agreement with previous studies linking *intI1* and *sul1* to anthropogenic activities in freshwater ecosystems [48, 61, 62]. Note that loading of traditional markers of water quality (total coliform and *E. coli*) also fell within the same quadrant as agricultural impacted sites. While the presence of these indicator organisms does not necessarily indicate the presence of harmful bacteria in water, their counts in APL and ADS may be heavily associated with agricultural run-off, effluents from human septic systems or from farm discharges as well as from non-point source fecal contamination such as wildlife or human recreational activities [63, 64].

**Figure 5.**
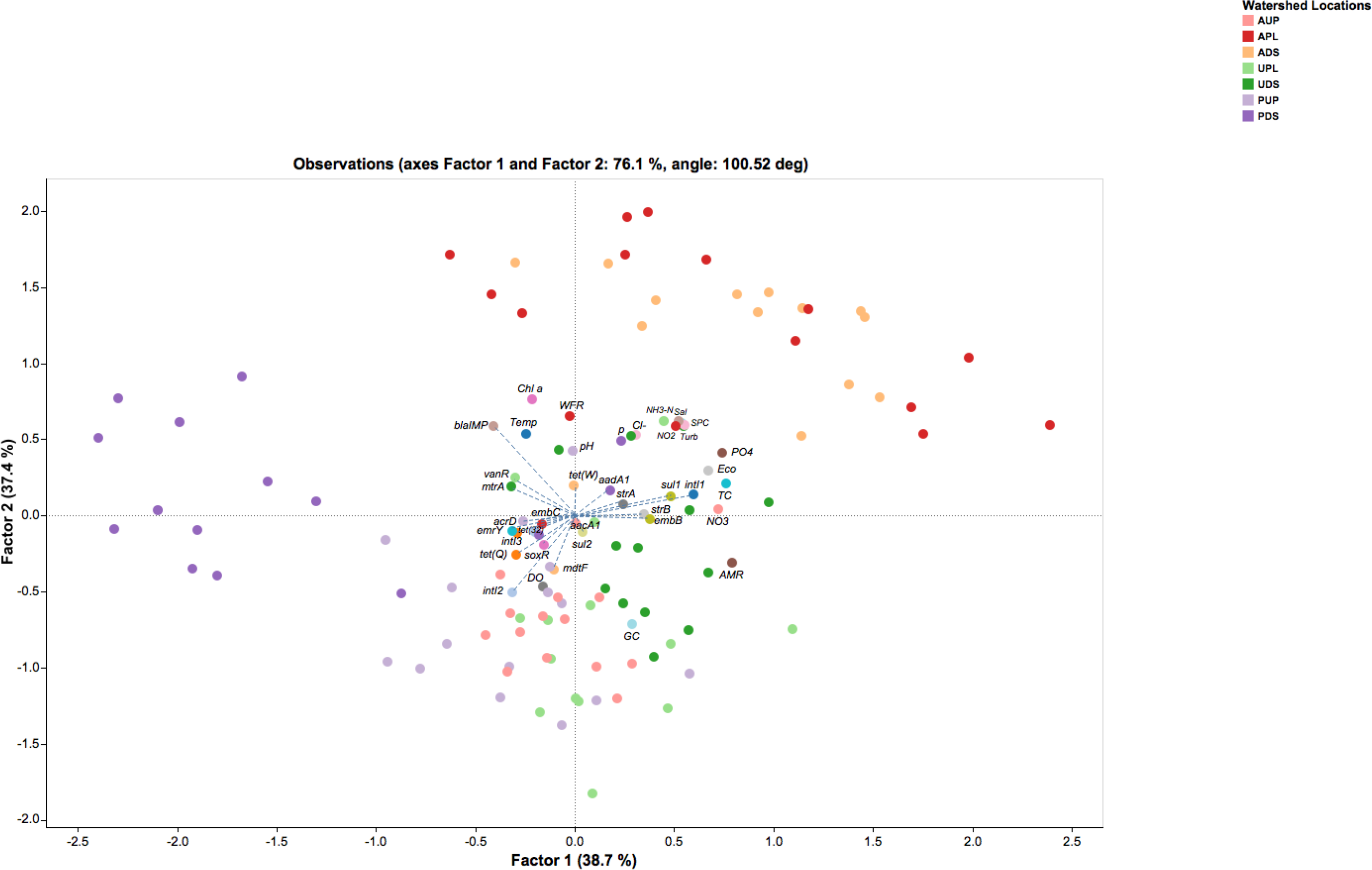
Factor analysis of antibiotic resistance genes/16S rRNA gene and environmental variables observed over time in watershed locations. AUP: agricultural upstream site; APL: agricultural polluted; ADS: agricultural downstream; UPL: urban polluted; UDS: urban downstream; PUP: protected upstream; PDS: protected downstream. Factor 1 and Factor 2 represent environmental stressors and aerobic conditions, respectively. Abbreviations: AMR: percentage of antibiotic resistance genes found in metagenomic sequences (based on CARD); Chl a: chlorophyll a; Cl^-^: dissolved chloride; DO: dissolved oxygen; Eco: *E. coli* counts; GC: percentage of guanine-cytosine content; NH_3_-N: ammonia; NO_2_: nitrites; NO_3_: nitrates; pH: potential of hydrogen; PO_4_: orthophosphates; Sal: salinity; SPC: specific conductivity; TC: total coliform counts; TDS: total dissolved solids; Temp: temperature; Turb: turbidity; WFR: water flow rate.

On the other hand, ratios of *emrY*, *mdtF*, *tet (Q)*, *soxR* appeared to be related to aerobic conditions as observed in AUP, UPL and PUP (Fig. 5). Significant positive correlations (p≤0.0284) were detected between this group of genes and dissolved oxygen (DO). These genes, associated primarily with aerobic and facultative anaerobic bacteria [65-67], may explain their higher relative abundance in the watersheds with higher DO levels. Whereas *intI2* was positively correlated with DO (r=0.4214, p<0.0001) but rarely detected in non-impacted sites (Fig. 4), we hypothesize that their relative abundance in watersheds aligns with the presence of aerobic bacterial hosts of integron genes rather than anthropogenic stressors or other environmental factors. Furthermore, integrase class 2 and *emrY*, *mdtF*, *tet (Q)*, *soxR* were positively correlated among them (p≤0.0103). It is possible that aerobic environments favor the ratio of these genes compared to the overall genes in a microbial community.

When comparing ratios of *acrD*, *embC*, *tet(32), intI3*, they were not affected by environmental stressors or aerobic conditions (Additional file 6: Figure S4a and Additional file 7: S4b). Instead, natural factors may influence their occurrence in the studied watersheds. For instance, *tet(32)*, previously documented in only anaerobic bacteria from clinical samples previously only documented in anaerobe bacteria from clinical samples [68, 69] has recently been reported to be widespread in human, animal and environmental resistomes [70]. Another example is the wide distribution of the lesser known integrase gene, *intI3*, in natural environments [71, 72]. The dynamics of *vanR* and *mtrA* genes, part of two-component systems conferring resistance to vancomycin and multiple antibiotics, respectively seemed also to be governed by the natural occurrence of bacteria [73, 74]. It is important to mention that the presence of biofilms in PDS may have also had an effect on the microbial community structure and thus of the naturally occurring ARGs. The role of biofilms as reservoirs of ARGs has been documented as has their influence on increasing resistance of bacteria to antibiotics 10-1000 times compared to free-living bacteria [75-77]. Finally, the ratio of *bla*_IMP_ seems as well to be aligned with the natural occurrence of bacteria in watersheds. This metallo-β-lactamase gene conferring resistance to carbapenems was positively correlated (p≤0.0233) with parameters such chlorophyll a and water flow rate (Figures 5, Additional file 5: Figure S3, Additional file 6: Figure S4a and Additional file 7: S4b). Although, an inverse relationship between chlorophyll a and water flow rate has been widely reported in freshwater ecosystems [78-82], this condition only applied in the more impaired watersheds (APL, ADS, UPL and UDS). In this study, we observed a positive correlation between both parameters in lesser-impacted environments such as AUP or non-impacted watersheds such as PUP. No association could be determined in PDS, perhaps due to the influence of pipe-associated biofilms on this site. Another correlation (p=0.0001) was observed between temperature and chlorophyll a, but this factor did not seem to have a major effect (p= 0.2733) on the population density harboring the *bla*_IMP_ gene. It should be noted that absolute GCNs of *bla*_IMP_ in urban, AUP and protected watersheds were detected in at least one order of magnitude higher than APL or ADS (Fig. 4 and Additional file 4: Figure S2). Additional analysis of phylogenetic information associated to *bla*_IMP_ genes (data not showed) suggested that species of *Pseudomonas* spp. are the most probable bacterial host for this gene. Members of this genus are important phytopathogens and are also opportunistic agents of human infections [83]. They have been reported in low nutrient or oligotrophic environments, urban environments, colonizing biofilms and plumbing structures [84]. These observations may explain the relative abundance of *bla*_IMP_ in non-agriculture impacted sites.

The model described in this study included only three of the major factors that may explain the overall variability of ARGs in watershed locations. Besides poor land-use practices, additional factors that may have influenced the pattern of bacterial community and ARGs in watersheds, such as seasonal conditions, water flow (i.e. flow in ADS is regulated by gated dams located 8.7 km further downstream), indirect human interventions (i.e. water collected in UDS passes through a 9.2 km pipeline) [57, 85].

### Analysis of antibiotic residues in freshwater samples

To further understand the impact of antibiotics on the aquatic environment, we screened for antibiotic residues using a subset of water samples (described in Methods section). The selection of antibiotic metabolites to screen for was derived from the metagenomics supported with information on the most commonly antibiotics employed for agricultural and human purposes in Canada [86-88]. Information from antibiotics used on farms was not available for this study area [89] compared to other studies [25, 90, 91]. Detection limits for the analytical methods used were relatively low for each antibiotic as follows: ampicillin (0.020 µg/L), sulfamethoxazole (0.0050 µg/L), chlortetracycline (0.025 µg/L), doxycycline (0.050 µg/L), oxytetracycline (0.010 µg/L) and tetracycline (0.025 µg/L). On analysis, none of the 3 groups of antibiotics were detectable in the subset of water samples. For various reasons, it is probable that the amounts of antimicrobials used in this part of Canada are a smaller fraction of those used in other countries (i.e. China) [92, 93]. Other factors such as a rapid degradation (including environmentally relevant conditions) [94-97], formation of metabolites (even at a higher concentration than the parent molecule) [98-101], and distribution (into sediments rather than the water column) [102-105], may have been resulted in these antibiotics not being detected. On the other hand, it is known that low concentrations of antibiotics or their metabolites have been associated with selectivity for antibiotic resistant bacteria [106, 107]. The observation of high richness of ARGs in the agriculture impacted sites compared to the urban (~1:7 ratio) and protected (~1:11 ratio) sites suggests that the ARGs were more likely derived from bacteria from the effluents rather than the de novo acquisition of resistance in the naturally occurring bacteria in the water. Finding greater GCNs of ARGs quantified in agriculture and urban impacted sites compared to non-impacted watershed locations, also supports this.

## Conclusions

The total number of sequences associated with ARGs from bacterial reads were low in aquatic environments (<1 %). The metagenomics approach identified a total of 78 different ARGs and one integrase class type 1 across all sampling sites. Agriculture impacted sites contained a higher richness of ARGs compared to the urban or protected environments. Nineteen genes were further screened to quantify GCNs of 1 ARG in the study sites. We also included integrase gene classes 2 and 3 due to their relevance in mobile genetic elements and ARGs. Using HT qPCR, we detected higher GCNs of ARGs in agriculture and urban impacted sites. A downstream transport pattern was identified for most of the ARGs during the dry season, while these differences became undetectable with the onset of higher precipitation in the study area. Genes such as *aacA1*, *aadA1*, *strA*, *strB*, *sul1*, *sul2* and *intI2* were more prevalent in agricultural sites, while that *tet(32)* and *tet(Q)* had a higher prevalence in the urban sites. Genes *tet(W)* and *intI1* were prevalent in both urban and agricultural settings. Moreover, an exploratory factor analysis found that there were three major contributors/drivers of ARGs in the watershed locations of this study: anthropogenic stressors (38.7 %), aerobic conditions (37.4 %) as well as natural occurrence (23.9 %). The inability to detect the antibiotics in the water suggests that the ARGs come from organisms from the effluents from the impacted sites. This is consistent with the resilience/stability of the antibiotic resistant organisms even after they enter the environment. Although the occurrence of ARGs in these sites was low compared to the bacterial population, high richness and copy numbers in agricultural sites and to a lesser extent in urban sites, demonstrates the influence of anthropogenic activities on the aquatic environment.

Abbreviations

ARGs: antibiotic resistance genes
GCNs: Gene copy numbers
HT qPCR: high throughput quantitative polymerase chain reaction
AUP: Agricultural upstream watershed sampling site
APL: Agricultural polluted watershed sampling site
ADS: Agricultural downstream watershed sampling site
UPL: Urban polluted watershed sampling site
UDS: urban downstream watershed sampling site
PUP: Protected upstream watershed sampling site
PDS: Protected downstream watershed sampling site

## Methods

### Sample collection

Forty liter samples were collected in sterile plastic carboys from three different British Columbia watersheds. A case-control design was used to characterize spatial and temporal distribution of representative antibiotic resistance and integrase genes in watersheds. Sampling within each site was conducted in 2-3 locations (upstream, at the ‘polluted’ site and downstream). Each sampling study site represented different land-use: agriculture (Agricultural Upstream or AUP, Agricultural Polluted or APL, and Agricultural Downstream or ADS); urban (Urban Polluted or UPL, and urban downstream or UDS); and non-impacted (Protected Upstream or PUP, and Protected Downstream or PDS). Descriptions of these sampling sites are detailed in Uyaguari et al. [53]. A total of 89 samples were collected within a one year period (April 2012-April 2013). Samples were pre-filtered *in situ* using a 105-µm spectra/mesh polypropylene filter (SpectrumLabs, Rancho Dominguez, CA), kept on ice, and then transported to the British Columbia Centre for Disease Control Public Health Laboratory for processing and storage at 4 °C (within 2-3 h). One 250-ml sample was collected in brown sterile bottles without prefiltration step transported to the laboratory, and stored at −80 °C for further analysis of antibiotics.

### Metadata

Water quality parameters were measured *in situ* using a YSI Professional Plus handheld multiparameter instrument (YSI Inc., Yellow Springs, OH). Physico-chemical parameters of watersheds included: temperature (°C), dissolved oxygen (mg/L), specific conductivity (µS/Cm), total dissolved solids (mg/L), salinity (PSU), pressure (mmHg), and pH. Turbidity (NTU) was measured using a VWR turbidity meter model No. 66120-200 (VWR, Radnor, PA). Water flow data (m^3^/s) was determined *in situ* using a Swoffer 3000 current meter (Swoffer Instruments, Seattle, WA). Total coliform and *Escherichia coli* counts were determined using the Colilert-24 testing procedure (IDEXX Laboratories, Westbrook, ME). Laboratory analysis included dissolved chloride (mg/L) and Ammonia (mg/L) using automated colourimetry (SM-4500-Cl-G) and spectrophotometry (SM-4500NH3G). Other nutrients such as chlorophyll a [108], orthophosphates [109], nitrites and nitrates [110] were also analyzed.

### Filtration and DNA extraction

The bacterial fraction of the water samples was captured by passing water through a 1 µm to remove protists and larger cells and 0.22-µm 142 mm Supor-200 membrane disc filters (Pall Corporation, Ann Harbor, MI) to capture bacterial cells, as described by Uyaguari-Diaz et al. [53, 111]. Bacterial-sized cells captured on the Supor-200 membrane disc filters were washed with 1X phosphate buffered saline (PBS) and 0.01 % Tween (pH 7.4). Mechanical procedures involving shaking, and centrifugation were used to remove and further concentrate cells (3,300 × g, 15 min at 4 °C). Cell aliquots of 300 µl were stored at −80 °C for further DNA extraction. DNA was extracted from multiple concentrated cell aliquots using the PowerLyzer PowerSoil DNA kit (MoBio, Carlsbad, CA) by following the manufacturer’s instructions. DNA was precipitated using 10 % 3M sodium acetate and 2X 100 % ethanol, and 5 µl of 5 µg/µl linear acrylamide, washed with 1 ml of 70 % ice-cold ethanol, and eluted in 34 µl of 10 mM Tris solution. DNA concentration and purity was assessed with Qubit fluorometer (Life Technologies, Carlsbad, CA) and NanoDrop spectrophotometer (NanoDrop technologies, Inc., Wilmington, DE), respectively.

### Metagenomic sequencing

To characterize microbial communities and subsequently identify antibiotic resistance elements in watersheds samples, libraries were prepared using Nextera XT DNA Sample Preparation kit (Illumina, Inc., San Diego, Ca).

Sequencing libraries were generated using 1 ng of DNA, according to the protocol described by the manufacturer’s instructions with a gel size selection modification [111]. Metagenomic sequencing was performed on a MiSeq platform (Illumina, Inc., San Diego, CA) using a MiSeq reagent kit V2 2x 250 bp paired-end output. Raw bacterial reads are available as part of a large-scale Watershed Metagenomics project, BioProject ID: 287840. (http://www.ncbi.nlm.nih.gov/bioproject/287840).

### Bioinformatics workflow

Trimmomatic version 0.32 [112] was used to remove adapters using the sequences packaged with the A5-Miseq assembly pipeline [113], discard low quality and short sequences (<75 nt). Sequences were then assembled using PandaSeq [114], unassembled pairs were also retained. De novo assembly was conducted on the assembled and unassembled pairs using MEGAHIT [115] and any contig shorter than 200 nt was discarded. Nucleotide sequences were aligned against the comprehensive antibiotic resistance database (CARD) [116] (available from http://arpcard.mcmaster.ca) and Integrall (available from http://integrall.bio.ua.pt) [117] using BLAST [118].

### Probe and primer design

Fully annotated genes from CARD encoding resistance to known antibiotics were used to design primer and probes for qPCR using Primer3Plus software [119]. Specific primers for *intI1-3* genes were selected from the literature [46]. Genes encoding for putative proteins such as elongation factors, alginates, or other hypothetical proteins were not considered for part of this analysis. Additional file 1: Table S2 summarizes primers and probes used in this study. All probes used a 5’ 6-FAM dye with an internal ZEN quencher and 3’ Iowa Black fluorescent quencher (Life Technologies, Carlsbad, CA).

### Standard curves

Genomic DNA from either agricultural or urban impacted watersheds was used as template to generate amplicons for each antibiotic resistance gene (Additional file 1: Table S2). Non-template controls used nuclease free-water (Promega Corporation, Fitchburg, WI). DNA from purified strains of *E. coli* ATCC 25922, and *E. coli* strains JVC1076, JVC1170, and JM109 (generous gift from Davies, Miao, and Villanueva, The University of British Columbia) were used as amplification controls for 16S rRNA gene, *intI1*, *intI2*, and *intI3* genes, respectively. PCR conditions were conducted as follows: 94 °C for 5 min, followed by 35 cycles of 94 °C for 30 s, 55 °C (50 °C) for 45s, 72 °C for 1 min and a final extension at 72 °C for 10 min. Amplicons were visualized on a 1.5 % agarose gel. PCR amplicons were purified with a QIAQuick PCR Purification Kit (Qiagen, Maryland, MD) according to the manufacturer’s instructions. The purified amplicons were ligated into pCR2.1-TOPO cloning vectors (Invitrogen, Carlsbad, CA) and transformed into One Shot *E. coli* DH5 α-T1^R^ competent cells following the manufacturer’s protocol. Transformants were grown overnight at 37 °C in lysogeny broth with 50 µg/ml of kanamycin. Plasmids were extracted and purified using PureLink Quick Plasmid Miniprep Kit (Life Technologies, Carlsbad, CA), and quantified using a Qubit dsDNA high sensitivity kit on a Qubit 3.0 fluorometer (Life Technologies, Carlsbad, CA). Plasmids for each antibiotic resistance gene and integrase gene class were end-sequenced using an ABI 3130xl Genetic Analyzer (Life Technologies, Carlsbad, CA) with M13 forward primer (-20) (5’-GTAAAACGACGGCCAG-3’) and M13 reverse primer (5'-CAGGAAACAGCTATGACC-3') using BigDye Terminator version 3.1 cycle sequencing kit (Applied Biosystems, Warrington, UK). The resultant set of DNA sequences for each antibiotic resistance gene and integrase classes were searched against the GenBank database using BLASTX with default settings.

Plasmid DNA harboring amplicon gene standards were linearized by digestion with the BamHI endonuclease (Life Technologies, Carlsbad, CA). Serial dilutions (n=6) of the linearized plasmids were multiplexed and used as templates to generate standard curves. Additional bioinformatics analysis screened for potential primer collision cases against Blast nt database using up to 1000 nt as maximum collision distance (Additional file 8). Estimates of bacteria, antibiotic resistance and integrase gene quantities in watershed sites were determined using 16S rRNA, ARGs and *intI* gene fragments, respectively (Additional file 1: Table S2). Gene copy numbers per ml of sample were calculated as previously described by Ritalahti et al. [51].

### High-throughput multiplex quantitative polymerase chain reaction

DNA extracts from watershed samples were diluted 10-fold and 1.25 µl of DNA from each sample was pre-amplified with low concentrated primer pairs (0.2 µM) corresponding to all assays (Additional file 1: Table S2 in a 5 µl reaction volume using TaqMan Preamp Master Mix (Life Technologies, Carlsbad, CA) according to the BioMark protocol (Fluidigm Corporation, South San Francisco, CA). Unincorporated primers were removed using ExoSAP-IT High-Throughput PCR Product Clean Up (MJS BioLynx Inc., Brockville, ON) and samples were diluted 1:5 in DNA Suspension Buffer (TEKnova, Hollister, CA).

The pre-amplified products were run on the BioMark system (Fluidigm Corporation, South San Francisco, CA) using 96.96 dynamic arrays. Five µl of 10x assay mix (9 µM primers and 2 µM probes) were loaded to assay inlet, while 5 µl of sample mix (2x TaqMan Mastermix (Life Technologies, Carlsbad, CA), 20x GE Sample Loading Reagent, nuclease-free water and 2.25 µl of pre-amplified DNA were loaded to each sample inlet of the array following manufacturer's recommendations. After mixing the assays and samples into the chip by an IFC controller HX (Fluidigm Corporation, South San Francisco, CA). Quantitative PCR was performed with the following conditions: 50 °C for 2 min, 95 °C for 10 min, followed by 40 cycles of 95 °C for 15s and 60 °C for 1 min. Samples were run in quadruplicates for all six standards, 89 environmental samples and one no-template control.

### Analysis of antibiotic residues in water

A subset of water samples stored at −80 °C were sent for analysis of antibiotic residues to a commercial laboratory. This subset included 13 water samples from APL and one sample of the other sites (AUP, ADS, UPL, UDS, PUP and PDS). Selection of antibiotic residues for testing was based on the major groups of antibiotic resistance genes found through the CARD database and included: ampicillin (β-lactam group); sulfamethoxazole (sulfonamide group); and chlortetracycline, doxycycline, oxytetracycline and tetracycline (tetracycline group). Residues of ampicillin and sulfamethoxazole were analyzed by LC/MS following the methods described in EPA 549.2 [120], while the tetracycline group was tested by LC/MS-MS following a modified EPA 549.2 method [120].

### Data analysis

Gene copy numbers and metadata were transformed using log_10_ function for analysis. Analysis of repeated measures was conducted on the qPCR data for ARGs and *intI* to detect differences among watershed sites over time. Replicate values of each antibiotic resistance and integrase genes per site were averaged and these values introduced into the model. Tukey’s test was used to determine time effect among the different sites. Spearman’s rank correlation analysis was also conducted among ARGs, *intI* and water quality parameters. Factor analysis employed a parsimax oblique rotation that included metadata and the ratio of each ARG normalized by the 16S rRNA gene (as estimated by HT qPCR platform) measured in watershed locations. Statistical analyses were performed using Statistical Analysis System (SAS, version 9.4 for Windows). A p-value of 0.05 was assumed for all tests as a minimum level of significance.

## Authors’ Contributions

MUD conceived, designed, performed the experiments and wrote the manuscript. MAC led the bioinformatics analyses and designed the experiments. ZL designed and performed the experiments. KC, MC, SL, WB performed the experiments.

MN performed size selection of sequencing libraries. DD and WH provided additional bioinformatics analyses. KM, JIR, PT and NP designed the experiments, contributed analysis tools, guided the analyses and aided in interpretations. All authors contributed to final revisions of the manuscript.

MN holds shares of Coastal Genomics, a privately owned British Columbia company offering the Ranger Technology used in this study. The authors declare that the research was conducted in the absence of any commercial or financial relationships that could be construed as a potential conflict of interest.

This work was funded by Genome BC and Genome Canada grant (LSARP-165WAT). MUD was supported by a Mitacs Accelerate Fellowship.

## Acknowledgements

The authors would like to thank Belinda Wong, Frankie Tsang and staff members at the BCCDC Public Health Laboratory, Environmental Microbiology Laboratory for their support. We also thank Brian Auk, Mark McCabe and staff in the Molecular Microbiology and Genomics Program at the BCCDC Public Health Laboratory for their sequencing expertise. We would like to thank Jared R. Slobodan (Coastal Genomics) for his assistance with gel size selection. Thanks to Dr. Robert Balshaw (UBC), David Andrade Laborde (Ryder System, Inc.) and Guillermo Baños Cruz (University of Guayaquil) for providing statistical advice. Finally, we would like to acknowledge Drs. Julian Davies, Vivian Miao and Ivan Villanueva at The University of British Columbia for providing integrase control strains.

